# Altered basal ganglia output during self-restraint

**DOI:** 10.1101/2022.04.23.489276

**Authors:** Bon-Mi Gu, Joshua D Berke

## Abstract

Suppressing actions is essential for flexible behavior. Multiple neural circuits involved in behavioral inhibition converge upon a key basal ganglia output nucleus, the substantia nigra pars reticulata (SNr). To examine how changes in basal ganglia output contribute to self-restraint, we recorded SNr neurons during a proactive behavioral inhibition task. Rats responded to *Go!* cues with rapid leftward or rightward movements, but also prepared to cancel one of these movement directions on trials when a *Stop!* cue might occur. This action restraint – visible as direction-selective slowing of reaction times – altered both rates and patterns of SNr spiking. Overall firing rate was elevated before the *Go!* cue, and this effect was driven by a subpopulation of direction-selective SNr neurons. In neural state space, this corresponded to a shift away from the restrained movement. SNr neurons also showed more variable inter-spike-intervals during proactive inhibition. This corresponded to more variable state-space trajectories, which may slow reaction times via reduced preparation to move. These findings open new perspectives on how basal ganglia dynamics contribute to movement preparation and cognitive control.

## Introduction

Fluid, efficient behavior often involves simply triggering well-learned behaviors. However, flexibility requires that such behaviors can be suppressed, should circumstances change. This capacity for behavioral inhibition is considered central to cognitive control (Bari and Robbins, 2013), and is compromised in a range of neurological and psychiatric disorders (Chambers *et al*., 2009).

Behavioral inhibition can be ‘reactive’ – for example, promptly responding to an unexpected *Stop!* cue by cancelling upcoming actions. Reactive inhibition has been shown to involve fast cue responses in frontal cortex and basal ganglia pathways (Jahanshahi *et al*., 2015; Wager *et al*., 2005), including from the subthalamic nucleus (STN) to SNr (Schmidt *et al*. 2013). This rapid response to stimuli can transiently, and broadly, retard action initiation, providing time for a second set of basal ganglia mechanisms to cancel actions (Mallet *et al*. 2016; Schmidt and Berke 2018).

By contrast, ‘proactive’ inhibition refers to a state of preparation, in which particular actions are restrained (Cai *et al*., 2011, Claffey *et al*., 2010). Proactive inhibition has been argued to be especially important for human life (Aron 2011), and is behaviorally apparent as longer reaction times (RTs) selectively for the restrained action. The underlying mechanisms are not well understood, but have been proposed (Aron 2011) to involve the pathway from striatum “indirectly” to SNr, via globus pallidus pars externa (GPe). In a prior study (Gu *et al*. 2020) we therefore recorded from GPe neurons as rats engaged proactive inhibition towards a specific action. The state of being prepared to stop did not involve an overall net change in GPe firing rate, but rather a shift at the level of neural population dynamics away from action initiation, and towards the alternative action. One objective of the present work was to determine whether a corresponding preparatory change in population dynamics is visible “downstream” in SNr, thereby altering basal ganglia output. Basal ganglia output neurons are thought to affect behavior not just via their firing rates, but also via their firing patterns and synchrony (Rubin *et al*., 2012). In particular, Parkinson’s disease (PD) is associated with an increase in firing variability and synchronous bursting, often without rate changes (Lobb, 2014; Willard *et al*., 2019). As PD is characterized by slowed movement initiation (Low *et al*., 2002), we assessed whether related physiological changes are present when movements are slowed as the result of proactive inhibition.

## Results

### Reaction times are selectively slowed with proactive inhibition

We used a selective proactive inhibition task (Gu *et al*., 2020), a variant of our extensively-characterized rat stop-signal task (Leventhal *et al*., 2012; Schmidt *et al*., 2013; Mallet *et al*., 2016) (Fig. 1A). Rats start a trial by nose-poking an illuminated start-port. To proceed, they need to maintain this position for a variable delay (500-1250ms, uniform distribution) until a *Go!* cue is presented (1 kHz or 4 kHz tone, indicating a leftward or rightward movement respectively). If the movement is initiated rapidly after the *Go!* cue (RT limit < 800ms), and completed correctly and promptly (movement time limit, MT < 500ms), rats are rewarded with a sugar pellet dispensed at a separate food hopper. On a subset of trials, the *Go!* cue is followed by a *Stop!* cue (white noise burst; delay after *Go!* cue onset = 100 - 250ms).This indicates that the rat should not initiate a movement, and instead hold their nose in the Center port for at least 800ms (after *Go!* cue onset) to trigger reward delivery.

**Figure 1.**
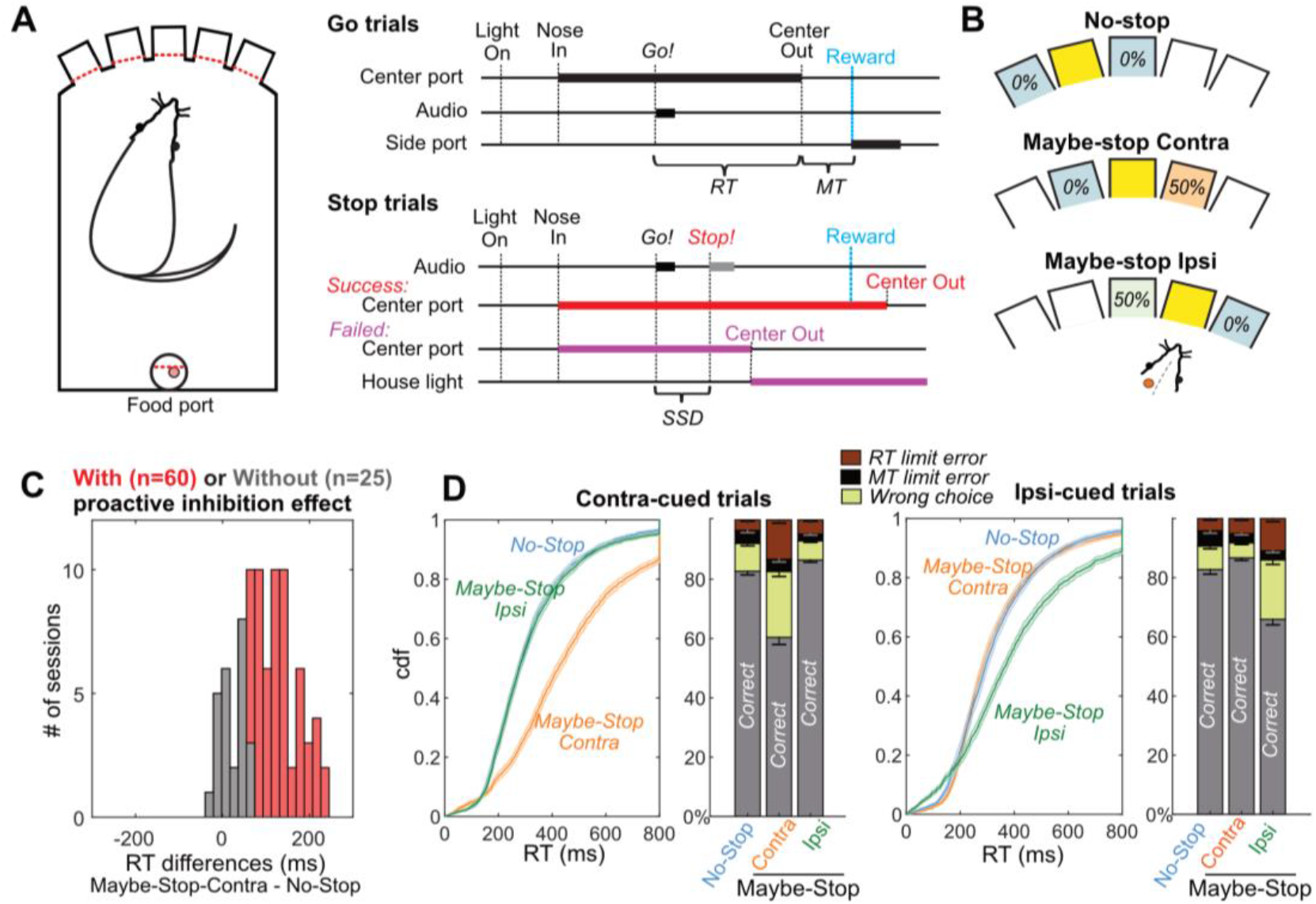
Selective proactive inhibition. (A) Left, operant box configuration, with dashed red lines indicating photobeams for nose detection; right, event sequence for Go and Stop trials. RT, reaction time; MT, movement time; SSD, stop-signal delay; Reward, delivery of a sugar pellet to the food port. (B) Trial start location indicates stop probabilities. In this example configuration with left SNr recording, illumination of the middle hole indicates that *Go!* cues instructing rightward movement may be followed by a *Stop!* cue, but *Go!* cues instructing leftward movements will not (“Maybe-Stop-Contra”). (C) Overall, the sessions in which SNr units were successfully recorded showed strong proactive inhibition effect (n=85, Wilcoxon signed rank tests on median RT differences between Maybe-Stop-Contra and No-Stop-Contra conditions, *p*=8.2e-15). Among them, the individual sessions are considered to show proactive inhibition (red) if the reaction time difference between Maybe-Stop-Contra and No-Stop trials is statistically significant (one-tail Wilcoxon rank sum test, *p*<0.05). (D) Cumulative distribution functions (cdf) of RTs of Maybe-Stop-Contra condition show selective slowing for the contra-cued trials, but not for the ipsi-cued trials. Response ratios also show selective increase of wrong choice and RT limit errors for the Maybe-Stop direction. Shaded band and error bars, S.E.M. across n = 60 sessions with proactive inhibition effect. RT limit error = nose remained in Center port for >800ms after *Go!* cue onset; MT limit error = movement time between Center Out and Side port entry >500ms.

To probe selective proactive inhibition, the three possible start ports were associated with different *Stop!* cue probabilities (counterbalanced across rats; Gu *et al*., 2020, Fig. 1B, Table 1). These were: no possibility of *Stop!* cue (“No-Stop”); 50% probability that a left *Go!* cue will be followed by the *Stop!* cue (“Maybe-Stop-left”); and 50% probability that a right *Go!* cue will be followed by the *Stop!* cue (“Maybe-Stop-right”). We obtained SNr microelectrode recordings from rats (n=10) that had successfully learned this proactive task, as indicated by significant and selective slowing of RTs for movements contraversive to the implant side (Fig. 1C). For example, if electrodes were placed in the right SNr, contraversive proactive inhibition would mean longer RTs for Maybe-Stop-left trials, compared to No-Stop trials. In the same rats we compared SNr activity in sessions in which this proactive effect was significant (n=60), to sessions in which it was not (n=25; Fig. 1C).

**Table 1.**
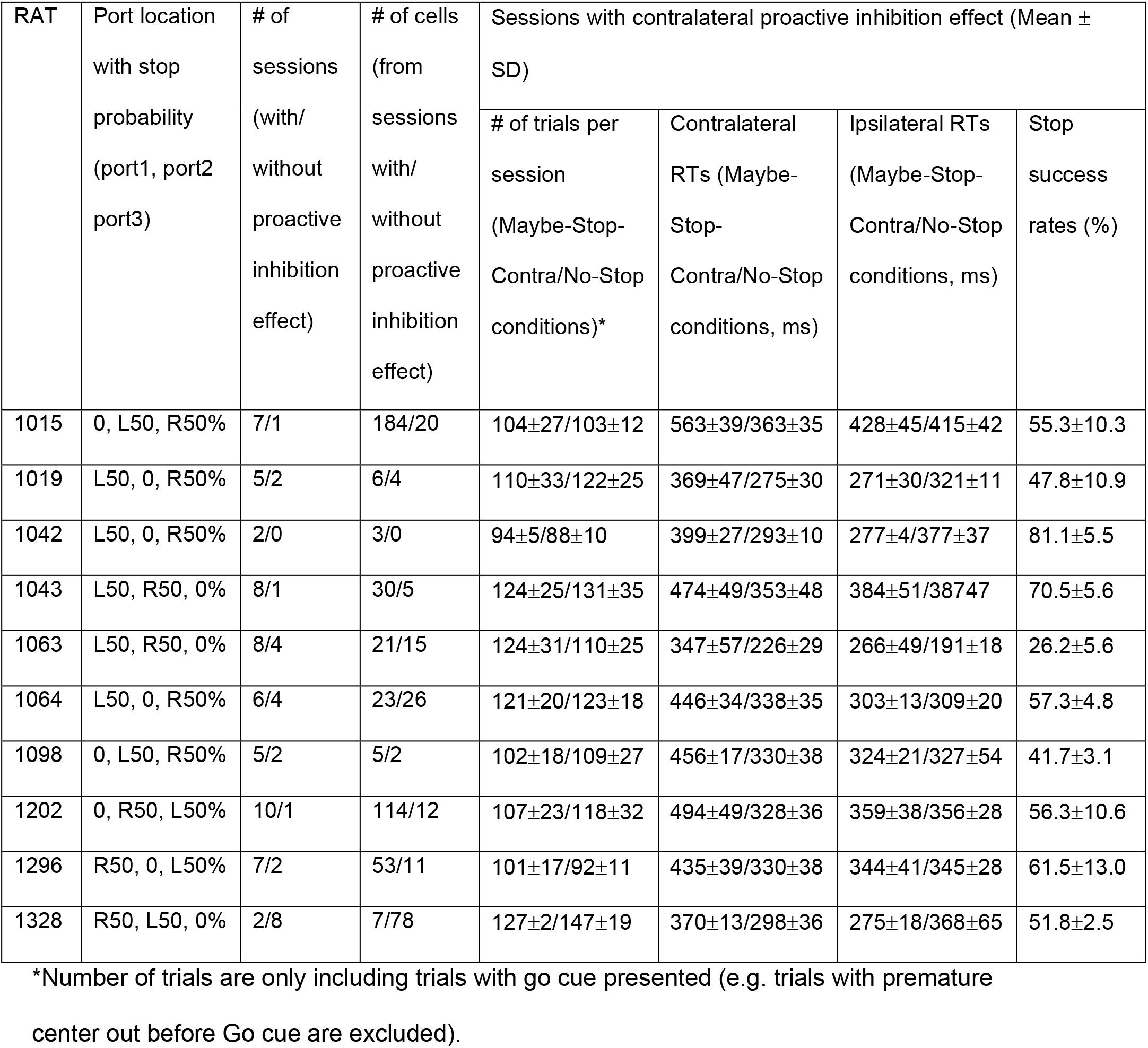
Information on individual rats.

| RAT | Port location<br>with stop<br>probability<br>(port1, port2<br>port3) | # of<br>sessions<br>(with/<br>without<br>proactive<br>inhibition<br>effect) | # of cells<br>(from<br>sessions<br>with/<br>without<br>proactive<br>inhibition<br>effect) | Sessions with contralateral proactive inhibition effect (Mean ± SD) | | | |
| --- | --- | --- | --- | --- | --- | --- | --- |
|  |  |  |  | # of trials per<br>session<br>(Maybe-Stop-Contra/No-Stop<br>conditions)* | Contralateral<br>RTs (Maybe-Stop-Contra/No-Stop<br>conditions, ms) | Ipsilateral RTs<br>(Maybe-Stop-Contra/No-Stop<br>conditions, ms) | Stop<br>success<br>rates (%) |
| 1015 | 0, L50, R50% | 7/1 | 184/20 | 104±27/103±12 | 563±39/363±35 | 428±45/415±42 | 55.3±10.3 |
| 1019 | L50, 0, R50% | 5/2 | 6/4 | 110±33/122±25 | 369±47/275±30 | 271±30/321±11 | 47.8±10.9 |
| 1042 | L50, 0, R50% | 2/0 | 3/0 | 94±5/88±10 | 399±27/293±10 | 277±4/377±37 | 81.1±5.5 |
| 1043 | L50, R50, 0% | 8/1 | 30/5 | 124±25/131±35 | 474±49/353±48 | 384±51/387±47 | 70.5±5.6 |
| 1063 | L50, R50, 0% | 8/4 | 21/15 | 124±31/110±25 | 347±57/226±29 | 266±49/191±18 | 26.2±5.6 |
| 1064 | L50, 0, R50% | 6/4 | 23/26 | 121±20/123±18 | 446±34/338±35 | 303±13/309±20 | 57.3±4.8 |
| 1098 | 0, L50, R50% | 5/2 | 5/2 | 102±18/109±27 | 456±17/330±38 | 324±21/327±54 | 41.7±3.1 |
| 1202 | 0, R50, L50% | 10/1 | 114/12 | 107±23/118±32 | 494±49/328±36 | 359±38/356±28 | 56.3±10.6 |
| 1296 | R50, 0, L50% | 7/2 | 53/11 | 101±17/92±11 | 435±39/330±38 | 344±41/345±28 | 61.5±13.0 |
| 1328 | R50, L50, 0% | 2/8 | 7/78 | 127±2/147±19 | 370±13/298±36 | 275±18/368±65 | 51.8±2.5 |

\*Number of trials are only including trials with go cue presented (e.g. trials with premature center out before Go cue are excluded).

The sessions with significant proactive inhibition effect show RT slowing selectively for the Maybe-Stop direction (Wilcoxon signed rank tests on median RTs of Maybe-Stop-Contra versus No-Stop: Contra cues: *z* = 6.8, *p* = 1.1e-11), but not for the other direction (Ipsi cues: *z* = -1.2, *p* = 0.22) (Fig. 1D). Additionally, on Maybe-Stop-Contra trials rats were more likely to fail to respond quickly enough (RT limit errors; Wilcoxon signed rank tests, *z* = 6.4, *p* = 1.5e-10) and to select the wrong choice (i.e. not matching the *Go!* cue; Wilcoxon signed rank tests, *z* = 6.5, *p* = 1.1e-10).

### Selective proactive inhibition recruits specific SNr subpopulations

We examined the task-related activity of individual SNr neurons (n=446; mean firing rate=38Hz, locations are shown in Supp. Fig. 1) recorded during the sessions with significant proactive inhibition. We first compared overall cell activity between Maybe-Stop-Contra and No-Stop trials. We focused on the epoch just before the *Go!* cue, as we presume that this time is critical for being “prepared-to-stop”. Average firing rates were significantly higher in the Maybe-Stop-Contra condition (Fig. 2A, top), and this difference was generated by a significant fraction of SNr neurons (Fig. 2A, bottom). Example cells show significant differences before the *Go!* cue between conditions (Fig. 2B). This elevated SNr firing with action restraint was not observed in the sessions without significant behavioral evidence for proactive inhibition (Fig. 2A, right; 173 neurons, mean FR=44Hz).

**Figure 2.**
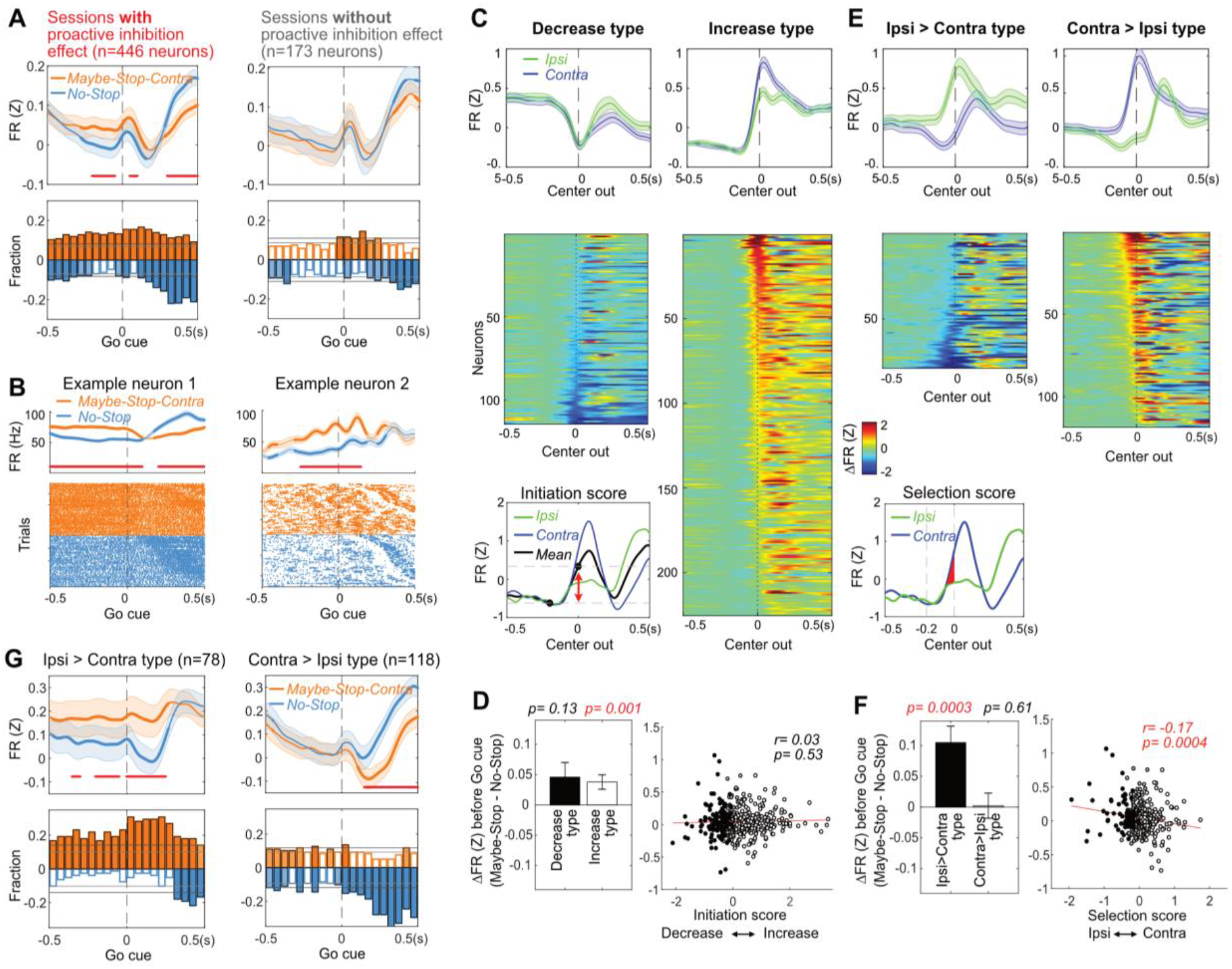
Elevated firing rates of specific SNr subpopulations with selective proactive inhibition. (A) Top left: before the Go! cue SNr firing is elevated in Maybe-Stop-Contra compared to No-Stop conditions. This occurs selectively in sessions with behavioral evidence of proactive inhibition (left). Each neuron’s firing rate is *Z*-scored and averaged (over all trials in which a *Go!* cue was presented, regardless of *Stop!* cues) Shaded band, +-S.E.M across n=446 (left) or n=173 neurons (right). Thicker lines indicate significant differences between conditions (*p*<0.05; Wilcoxon signed rank tests at each time point) and red lines at the bottom indicate times with significant difference remaining after Bonferroni correction (by the number of 50ms time bins; *p*<0.05). Bottom left: fraction of SNr neurons whose firing rate significantly differs between conditions, across time (*p*<0.05; Wilcoxon rank sum tests in each 50ms bin). Higher firing with Maybe-Stop-Contra, No-Stop conditions are shown as positive (orange) or negative (blue), respectively. Horizontal grey lines indicate thresholds for a significant proportion of neurons (binomial test, *p*<0.05 without or with multiple-comparisons correction, light and dark grey lines respectively). Light and dark color-filled bars are those for which the threshold was crossed without or with multiple-comparisons correction. Right, sessions without significant behavioral evidence of proactive inhibition do not show this firing rate difference between conditions (same format as left panels; n=173 neurons). (B) Two individual example neurons demonstrating the proactive elevation of firing rate before the *Go!* cue. Top: averaged firing rates in each condition. Shaded band, +-S.E.M across trials. Bottom: raster plots of individual trials. Trials are sorted by RTs. (C) Neurons were categorized as decrease-type or increase-type, based on an “Initiation Score”. We defined the “Initiation Score” for each neuron as the change in (*Z*-scored) firing rate in the 0.2s before Center Out (inset shows example neuron). Plots show average firing of each subpopulation (top; +-S.E.M) and individual cells sorted by Initiation Score (bottom). (D) No relation between Initiation Score and proactive inhibition (assessed as the difference between Maybe-Stop-Contra and No-Stop trials, in the 200ms before *Go!* cue). Bar graph (inset) shows that on average, both increase- and decrease-type neurons modestly increase firing with proactive inhibition (Wilcoxon signed rank tests in each group). Error bar is +-S.E.M across neurons. (E) We defined the “Selection Score” for each neuron as the integral of the difference in (*Z*-scored) firing rate between Contra and Ipsi actions during the 0.2s epoch before Center Out. Remainder of panel is as C, but for Selection Score.(F) Significant negative correlation between Selection Score and proactive inhibition. Bar graph (inset) shows that Ipsi>Contra neurons preferentially increase activity on Maybe-Stop-Contra trials. (G) Same result as F, using the format of panel A to illustrate time course.

A common categorization of SNr neurons distinguishes those that increase, versus decrease, firing rate in conjunction with behavioral events (e.g. Bryden *et al*., 2011; Fan *et al*., 2012; Gulley *et al*., 2002; Sato and Hikosaka, 2002). Decreases in firing shortly before movement onset are thought to enable movements, by disinhibiting downstream structures including the superior colliculus (Hikosaka and Wurtz, 1983b). If “decrease-type” neurons are receiving more excitation during proactive inhibition, this might delay their decrease to the level needed to release movements, resulting in longer reaction times. We therefore hypothesized that proactive inhibition is associated with elevated firing specifically of decrease-type neurons.

We categorized cells as increase-type or decrease-type based on their change in firing rate during the 200ms preceding movement onset (Fig. 2C). Increase-type were more numerous, as previously reported (Bryden *et al*., 2011; Joshua *et al*., 2009). Contrary to our hypothesis, both increase-type and decrease-type neurons contributed to elevated SNr activity with action restraint (Fig. 2D, left). There was no relationship between the extent of elevated firing in Maybe-Stop trials, and the firing rate change before movement onset (Fig. 2D, right).

We then examined whether proactive inhibition effects were related to neurons’ response selectivity, operationally defined as the firing rate difference between Contra and Ipsi actions just before Center Out (Fig. 2E, inset). We found a significant relationship: neurons more active just before Ipsi compared to just before Contra movements (“Ipsi > Contra”) showed elevated firing when Contra actions might need to be cancelled (Fig. 2F,G). No such relationship was found for neurons with the opposite selectivity (“Contra > Ipsi”; Fig. 2F,G).

We next considered the interaction between direction selectivity and increases vs. decreases in firing, in proactive inhibition. A cell could be classified as “Ipsi > Contra” because it preferentially increases firing with Ipsi movements, or because it preferentially decreases firing with Contra movements. We found that both subtypes had elevated firing before the *Go!* cue on Maybe-Stop-Contra trials (Supp. Fig. 2). Therefore, the proactive effect was not simply a matter of SNr cells that pause with Contra movements starting from a higher baseline rate, though this may contribute.

### Restraining one action biases population dynamics towards the alternative action

The elevated average firing rate of Ipsi > Contra cells suggests a preparatory bias towards Ipsi action, at times when Contra actions might need to be cancelled. To examine this further we turned to a state-space analysis, as we previously used with GPe data (Gu et al., 2020). We extracted principal components from the average firing rates of each neuron during Contra and Ipsi movements (Supp. Fig. 3) and used these to visualize neural population trajectories (Fig. 3A, top). We defined “Initiation” and “Selection” axes as the common, and distinct, aspects of these trajectories respectively, during the 200ms before action initiation. In particular, the Initiation Axis is a line drawn between the average state space positions at 200ms and 0ms relative to Center Out (disregarding the direction of movement). The Selection Axis connects the midpoints of the Contra and Ipsi trajectories (averaging across the same time epoch).

**Figure 3.**
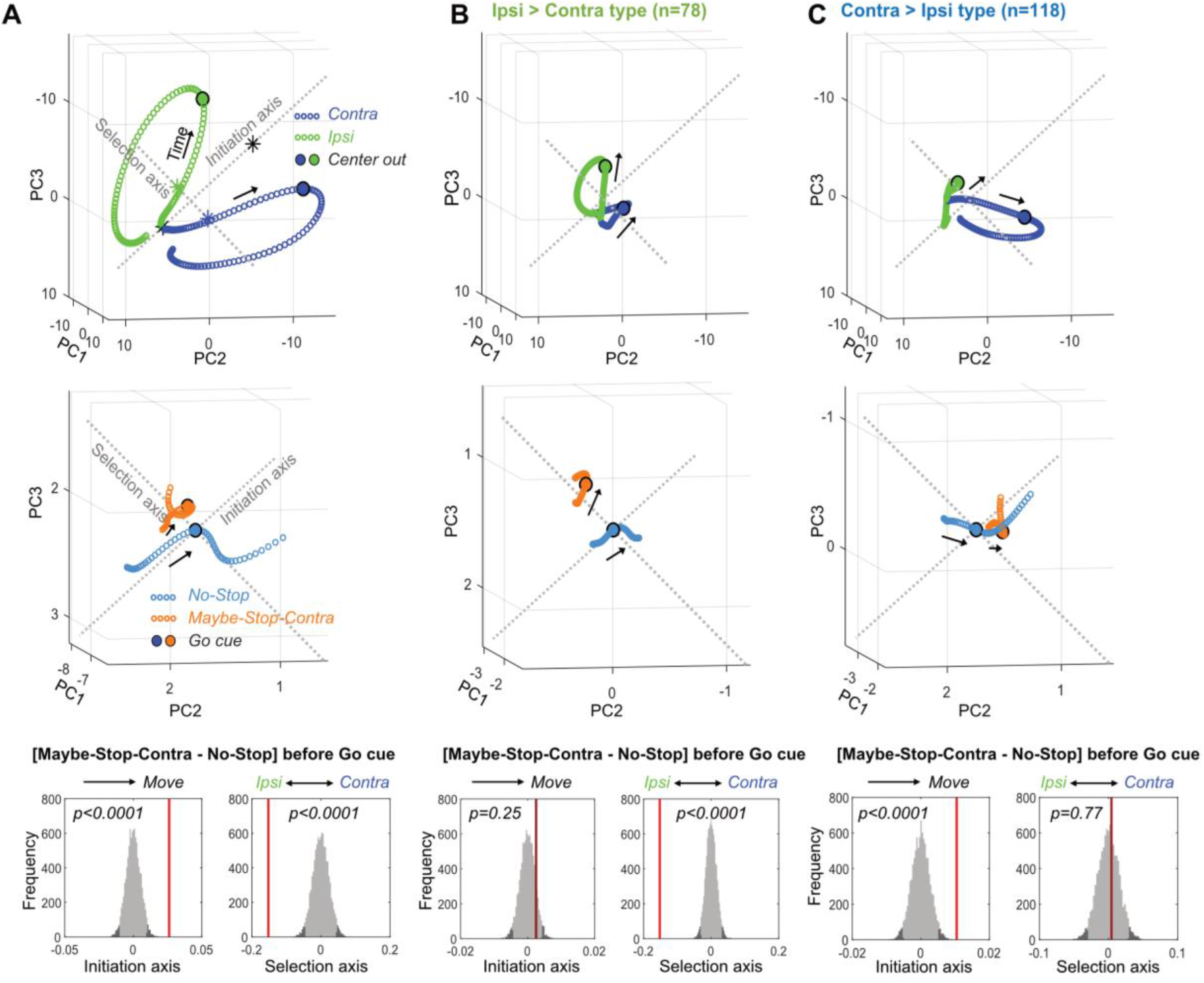
Biased SNr population dynamics during proactive inhibition. (A). Top, Overall SNr state-space trajectories before and during Contra (blue) and Ipsi movements (green), in the state space of the first 3 principal components (PCs). Trajectories show +-250ms around detected movement onset (Center Out, larger circles), with each small circle separated by 4ms. ‘Initiation Axis’ joins the positions (black asterisks) 200ms before and at action initiation (averaging Contra and Ipsi actions). ‘Selection Axis’ joins the means of each trajectory in the same epoch (colored asterisks). Middle, Comparing Maybe-Stop-Contra (orange) and No-Stop (blue) trials (+-100ms around Go! cue) in the same state space as above. Overall population state is visibly biased toward Ipsi along the Selection Axis. Bottom, Permutation tests of bias along each axis (average during -200ms to 0 relative to *Go!* cue), using all 10 PCs. Red bars, observed results; Grey, distributions of surrogate data from 10000 random shuffles of trial type labels. Dark grey indicates 5% of distributions at each tail. (B). As A, but for Ipsi > Contra cells only. The proactive bias towards Ipsi along the Selection Axis is more clearly visible. For comparisons, the PC dimension scale and Initiation/Selection axis are matched to the graph in A. (C). As A-B, but for Contra > Ipsi cells. These do not show a proactive bias on the Selection Axis, but on the Initiation Axis instead.

When we projected Maybe-Stop-Contra data into this space, we found that overall SNr population activity showed a clear shift before the *Go!* cue (Fig. 3A, middle). This shift was especially pronounced on the Selection Axis, with a highly significant bias towards Ipsi (Fig. 3A, bottom). There was also a significant bias on the Initiation Axis (towards movement). These two biases were associated with distinct functional cell classes (Fig. 3B, C). Ipsi > Contra cells showed a strong Selection Axis bias, without a significant Initiation Axis bias (Fig. 3B), and the converse was seen for Contra > Ipsi cells (Fig. 3C). This finding provides further evidence that Ipsi > Contra cells generate an Ipsi bias during selective proactive inhibition.

Moreover, the state of preparation was related to the subsequent outcome of the trial. Trajectories during wrong choice trials were biased both toward Ipsi action and towards initiation at the time of *Go!* cue (Supp. Fig. 3D), consistent with our prior GPe results (Gu *et al*., 2020).Trials in which rats failed to initiate actions (limited hold violations) did not show any statistically significant bias at the *Go!* cue time, although after the *Go!* cue the trajectory showed movement away from action initiation (Supp. Fig. 3D).

### Less regular firing with proactive inhibition

As movement slowing has been associated with changes in basal ganglia firing patterns in Parkinson’s Disease (Sharott *et al*., 2014; Tai 2022), we examined whether changes in SNr firing patterns accompany slower movement initiation during proactive inhibition. For each neuron we calculated the coefficient of variation (CV) of inter-spike intervals, and the proportion of spikes within bursts (Fig. 4A, B; using the Poisson surprise method; Legendy and Salcman 1985). To obtain sufficient numbers of inter-spike-intervals, both analyses used longer (3s) epochs before the *Go!* cue.

**Figure 4.**
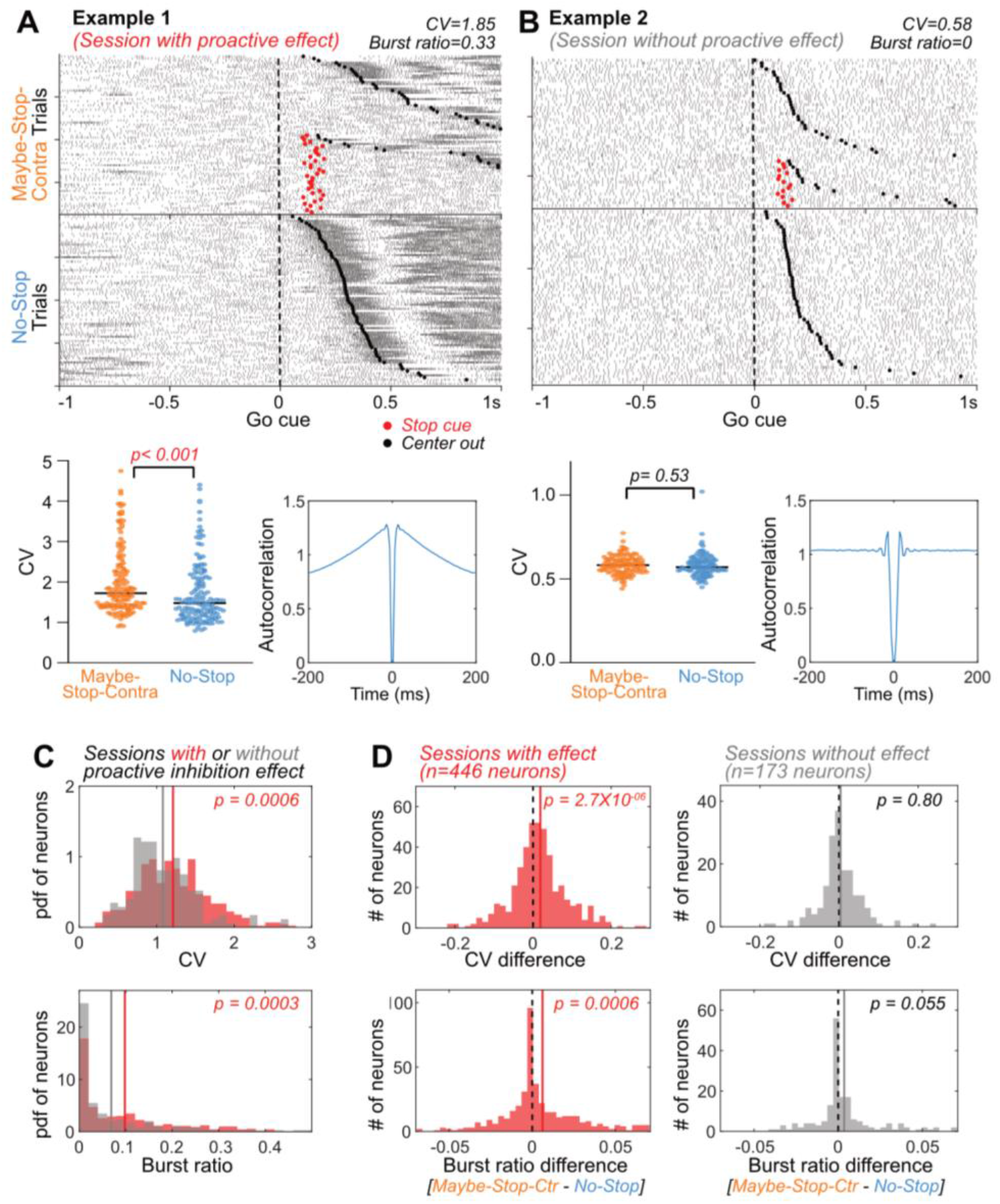
SNr firing is more irregular and bursty with proactive inhibition. (A) An example neuron from a session with proactive inhibition effect, showing spike rasters (Top; aligned on contra *Go!* cues, sorted by RTs), CVs of individual trials (Bottom left, for the 3s preceding Go*!* cues, Wilcoxon rank sum test) and autocorrelograms (Bottom right; for the 3s preceding *Go!* cues). CV: coefficient of variation of inter-spike intervals. (B) An example neuron from a session without a proactive inhibition effect. Same format as in (A). (C) Elevated CV and burst ratio of neurons recorded in sessions with behavioral evidence of proactive inhibition, compared to sessions without (Wilcoxon rank sum tests). This effect was also seen at the level of individual rats (Supp. Fig. 4C). Grey and red lines indicate mean of sessions with and without proactive inhibition effect, respectively. (D) Within-session comparison of Maybe-Stop-Contra and No-Stop trials shows increased CV and burst ratio when proactive inhibition is engaged (left). This effect is not present on sessions without significant proactive slowing of reaction times (right). Wilcoxon signed rank tests. Dotted black lines indicates zero and colored lines indicate mean of the neurons.

Proactive slowing was associated with altered spiking variability, in several distinct analyses. Neurons recorded in sessions with significant proactive inhibition behavior (n=446) showed more irregular and bursty firing compared to neurons (n=173) in sessions that did not (Fig. 4C). The degree of irregularity was correlated with the mean session-wide reaction times (Supp. Fig. 4A). Increased irregularity and bursting were also seen during Maybe-Stop-Contra trials compared to No-Stop trials, selectively in those sessions with significant proactive inhibition effect (Fig. 4D, left), and not those without (Fig. 4D, right). Moreover, the degree of increased irregularity between conditions was correlated with the magnitude of the proactive effect on reaction times (Supp. Fig. 4B).

### Proactive inhibition increases the variability of neural trajectories

How could increased spiking variability contribute to the slowing of RTs? At the population level, spike variability would correspond to more erratic state-space trajectories before the *Go!* cue (Fig. 5A). This would result in a more variable state at the (unpredictable) time of *Go!* cue onset. If effective movement preparation involves positioning neural activity within a “optimal subspace”, as previously proposed (Churchland *et al*. 2006), this more variable state would in turn result in RTs that are longer (on average) and more variable (across trials).

**Figure 5.**
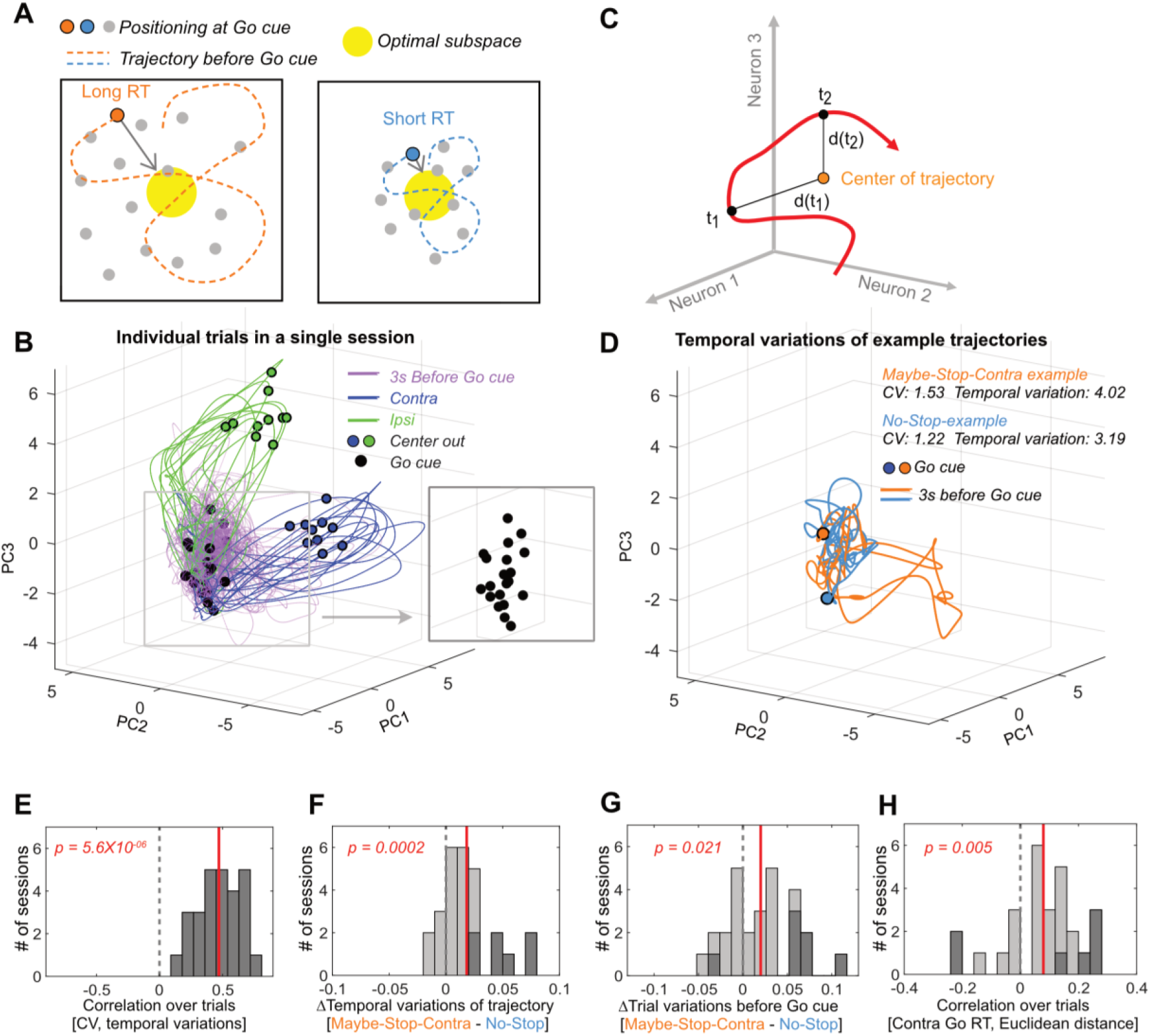
Altered variability of state trajectories with proactive inhibition. (A) Conceptual illustration for the relationship between trajectory variability and RTs. Larger fluctuations in trajectory will tend to result in a position further away from the “optimal subspace” when the *Go!* cue arrives. (B) Individual trial trajectories (10 trials each for Contra and Ipsi movements) for one example session (n=46 neurons). Trajectories are shown after PCA for visualization. (C) Trajectory variability was defined as the mean of the Euclidean distances at each time point to the mean position over the trajectory (in the full neural state space, without PCA). (D) Example trajectories for Maybe-Stop-Contra (orange) and No-Stop (blue) trials, before the *Go!* cue. Trajectories are shown after PCA for visualization. (E) Correlations between CV and trajectory variability for each session. (F) Trajectory variability increased on Maybe-Stop-Contra, compared to No-Stop trials. (G) Across trials, the state space position at *Go!* cue (−200ms to 0) was more variable for Maybe-Stop-Contra, compared to No-Stop trials. (H) Variability across trials of the state space position at Go! cue (−200ms to 0) was positively correlated with RT. Trials with less than 100ms reaction times were excluded because they would have already initiated the movement trajectory. For (E)-(H), sessions with more than 5 neurons (# of session = 27) were used for analysis (Wilcoxon signed rank test across session values). Dotted grey line indicates zero and red line indicates mean of the session values. Dark grey bars indicate sessions showing significant correlation (*p*<0.05 for (E), (H)), or conditional differences (Wilcoxon rank sum test, *p*<0.05 for (F), (G)).

To assess this idea we quantified trajectory fluctuations on individual trials (Fig. 5B-D). We included sessions (n=27) in which more than 5 neurons were recorded simultaneously, and examined trajectories in the 3s window before the *Go!* cue (for comparison to our CV measure). In the same way as CV measures variability over time of an individual neuron (one dimension) compared to its mean rate, we can measure the within-trial variability of a population (*n* dimensions, without PCA) by comparing the Euclidean distance of each time point to the mean position (Fig. 5C).

As expected, there was a strong relationship between CV of individual neurons and their corresponding population trajectory variability (Fig. 5E). Furthermore, trajectory variability was significantly increased by proactive inhibition (Figs. 5F). Trajectory variability increased during the hold period (examining the 0.8s epoch just before the *Go!* cue, excluding trials with hold duration <0.8s, Wilcoxon signed rank test, *p*=0.003), and was also observed even beforehand (0.8s epoch before Nose In, Wilcoxon signed rank test, *p*=0.003). This increase in within-trial trajectory variability with proactive inhibition indeed resulted in a more variable state space position at the time of *Go!* cue onset, across trials (Fig. 5G). Consistent with our hypothesis, this variability of state-space position across trials was correlated with RT (Fig. 5H).

### Proactive modulation of firing rate, and variability, are dissociable

Changes in preparation to move or stop can be evoked by explicit cues, as in our task design, but also by the subject’s ongoing experience – notably, what happened on the previous trial (Bissett and Logan, 2011; Pouget *et al*. 2011). We therefore examined how such ongoing experience affects the proactive influences over SNr firing. Regardless of previous trial type Maybe-Stop-Contra trials had slower RT (Fig. 6, top row), and an Ipsi-biased SNr state-space position (Fig. 6, third row). By contrast, the increased CV in SNr spiking was particularly apparent following Stop-fail trials (Fig. 6, bottom). This suggests that increased spike variability occurs when rats are especially concerned to avoid hasty responses, having just failed to sufficiently restrain behavior on the previous trial.

**Figure 6.**
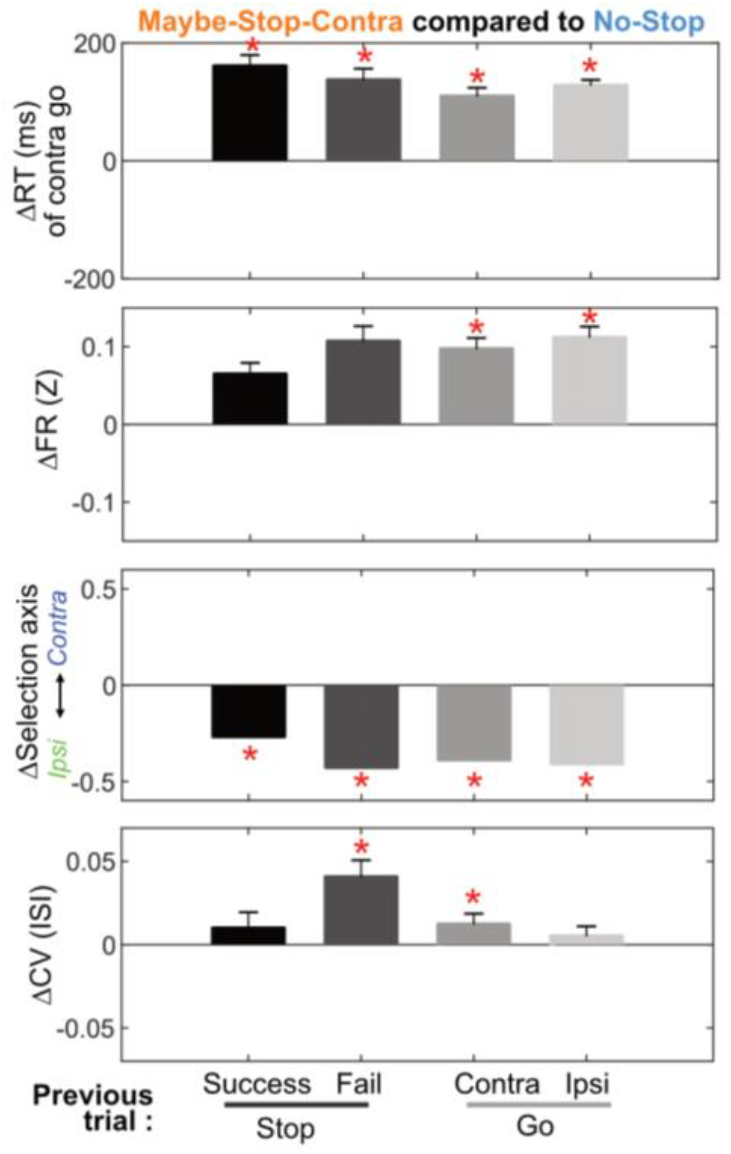
Feedback effect by previous trials and relation to trial outcomes. Differences in reaction time, firing rates, selection axis positions, and CVs of Maybe-Stop-Contra trials, compared to the No-stop condition (zero), and separated by previous trial type. The effect of previous trial type reached significance for the difference in CV (Friedman’s test, *X*^*2*^(3)=13.38, *p*=0.004), but not for contra go RT (*X*^*2*^(3)= 2.84, *p*=0.42) nor in firing rates (*X*^*2*^(3)= 5.09, *p*=0.17). Firing rates and selection axis measures use Ipsi > contra cells (during 200ms before *Go!* cue); all cells are included in the CV calculation. \**p* <0.05 with Bonferroni multiple comparison correction (Compared to No-Stop trials, permutation test as in Fig 3 for Selection axis, and Wilcoxon signed rank test for others). Error bars are +-S.E.M across sessions (n=60) for reaction times and across neurons (n=446) for firing rates and CV.

## Discussion

Our capacity to inhibit actions and thoughts can be influenced by a wide range of factors (Bari and Robbins 2013). These include external cues (such as warning stimuli) and “internal” processes such as attention and motivation (Meyer and Bucci 2016). Our experimental design provides some useful constraints on which factors are relevant to our results. We used a standard, operational definition of selective proactive inhibition: slowing of RTs for the particular movement that *might* need to be cancelled. Since this effect was observed even when no *Stop!* cue was actually presented, and was direction-selective, it cannot be explained simply by (for example) priming of the more global, cue-evoked Stop mechanisms that support reactive inhibition (i.e. “preparation to stop”). Instead, it implies an altered state already present by the time of the *Go!* cue.

We found evidence for this altered state in two aspects of SNr spiking. A subpopulation of movement-selective SNr neurons showed elevated firing before the *Go!* cue, and this was associated with a shift in population activity away from the specific restrained action and towards the alternative. At the same time, more erratic SNr firing results in a more variable state at the time of the *Go!* cue. This variability is associated with slowed RTs, plausibly because effective movement preparation involves achieving a more constrained range of network activity. Moreover, these firing rate and variability modulations reflect two dissociable mechanisms for proactive control, as they were recruited differently depending on events on the prior trial.

### Basal ganglia dynamics and the nature of restraint

Our results add to prior evidence that, beyond simple sensory or motor correlates, SNr firing is modulated by internal factors such as task context (Hikosaka and Wurtz 1983a; Lintz and Felsen 2016). At the same time our operational definition of proactive inhibition does not fully constrain which internal factors are at play – and these may vary both within, and between, individual subjects. Basal ganglia activity is especially affected by changes in reward expectation (Lauwereyns *et al*. 2002) including in SNr (Sato and Hikosaka 2002; Yasuda and Hikosaka 2017), producing faster RTs towards more rewarding movements. In our proactive task the Maybe-Stop direction receives rewards at a lower rate (due to failures-to-Stop), so asymmetrical reward expectation might be at least partially responsible for the neural and choice bias towards Ipsi movements on Maybe-Stop-Contra trials. However, the slower Contra RTs on Maybe-Stop-Contra trials were not accompanied by faster Ipsi RTs, suggesting that slowing of Contra movements did not result simply from a greater preparation of Ipsi movements.

The SNr state-space bias during proactive inhibition may arise from the “indirect pathway” projection from GPe, which we found to have a similar neural bias towards Ipsi movements in a prior study (Gu *et al*., 2020). However, proactive inhibition produced no overall change in GPe firing rates, and we were not able to identify a distinct subpopulation of modulated GPe neurons. The reasons for this GPe : SNr difference are not clear. Recent modeling has demonstrated that the influence of the GABAergic GPe inputs over SNr neurons can be far more complex (Simmons *et al*. 2020) than shown in classic rate models of basal ganglia function, for example switching from inhibitory to excitatory in an activity-dependent manner (Phillips *et al*. 2020). Alternatively, the shift in SNr Ipsi > Contra neuron firing with proactive inhibition may reflect other basal ganglia inputs, particularly the direct pathway input from striatum, or hyperdirect via the subthalamic nucleus (Schmidt *et al*. 2013).

### Behavioral control and neural variability

Our study also leaves unresolved the origins of the increased SNr spike variability we observed with proactive inhibition. Like GPe and STN neurons, SNr neurons are intrinsic pacemakers that spontaneously fire regularly even if their inputs are blocked (Zhou and Lee 2011). These inputs – which include the aforementioned direct and indirect pathways, and also extensive local collaterals within SNr (Brown *et al*., 2014; Mailly *et al*., 2003) – thus alter SNr spiking by either accelerating or delaying the occurrence of the next spike. Changes in spike time variability presumably reflect either changes in local network properties, e.g. due to neuromodulation (Delaville *et al*. 2012) or changes in the statistics of extrinsic inputs.

More variable SNr inter-spike-intervals within trials were associated with more variable trajectories through state-space, a more variable network state at *Go!* cue, and longer and more variable RTs. In the cortex, decreases in spike variability have been previously linked to various processes including stimulus onset (Churchland *et al*., 2010), attention (Cohen and Maunsell, 2009), and movement preparation (Churchland *et al*., 2006). There is also evidence of within-trial changes in spike variability in the striatum (Berke 2011). To our knowledge, there are no prior observations of specific task cues evoking *increases* in variability between or within trials. Our finding that variability increases with proactive inhibition – in particular following Stop-fail trials – suggests that variability might be actively elevated as part of a behavioral strategy. This would fit with proposals that neural variability can confer behavioral advantages, such as increased flexibility (Waschke *et al*. 2021). Alternatively, increased neural variability with proactive inhibition may simply reflect the *absence* of a reduction in variability that accompanies movement preparation. In this way, preparation to stop would consist, at least in part, of less preparation to go. Either way, our results demonstrate that cognitive control strategies can operate through shifts in neural variability.

Connecting single-neuron measures of variability to state-space concepts of movement preparation offers an intriguing perspective on PD, which is characterized by both slowed movements (Low *et al*., 2002) and elevated single-neuron spike variability (Dorval *et al*., 2008; Lobb, 2014; Willard *et al*., 2019). A more variable state of preparation may help explain why *average* RTs and movement times are slowed in PD, yet the fastest movements are still preserved (Mazzoni *et al*. 2007). An important goal for future studies is to better understanding how shifts in neural variability occur in the basal ganglia, and whether they originate from the same mechanisms during both proactive inhibition and Parkinson’s Disease.

## Acknowledgements

We thank Charles Wilson and members of the Berke Lab for valuable feedback. This work was supported by UCSF, NIH (R01MH101697, R01NS123516), and CHDI.

## Author contributions

B.G., Conceptualization, Data curation, Software, Formal analysis, Investigation, Visualization, Methodology, Writing – original draft; J.D.B., Conceptualization, Resources, Supervision, Funding acquisition, Writing – original draft, Writing – review and editing.

## Materials and methods

### Animals

All animal experiments were approved by the University of California, San Francisco Committee for the Use and Care of Animals. Adult male Long-Evans rats were housed on a 12 hr/12 hr reverse light-dark cycle, with training and testing performed during the dark phase.

### Behavior

The rat proactive stop task has been previously described in detail (Gu *et al*., 2020). Briefly, rats were trained in an operant chamber (Med Associates, Fairfax VT) which had five nose-poke holes on one wall, a food dispenser on the opposite wall, and a speaker located above the food port. Each trial starts with one of the three starting ports illuminated to indicate the *Stop!* cue probabilities (‘Light On’), and the same start port was repeated for 10-15 trials. Inter-trial intervals were randomly selected between 5 and 7s. The mapping of stop probabilities to nose-poke locations are counter-balanced between rats but maintained in each rat (Table 1).

### Electrophysiology

We recorded SNr data from ten rats (all animals in which we successfully recorded SNr neurons during contraversive proactive inhibition). Each rat was implanted with 15 or 30 tetrodes in bundles bilaterally targeting SNr (Supp. Fig1). Wide-band (0.1–9000 Hz) electrophysiological data were recorded with a sampling rate of 30000/s using an Intan RHD2000 recording system (Intan Technologies). All signals were initially referenced to a skull screw (tip-flattened) on the midline 1 mm posterior to lambda. For spike detection we re-referenced to an electrode common average, and wavelet-filtered (Wiltschko *et al*., 2008) before thresholding. For spike sorting we performed automatic clustering units using MountainSort (Chung *et al*., 2017) followed by manual curation of clusters. Approximately every 2-3 sessions, screws were turned to lower tetrodes by 100-160 µm; to avoid duplicate neurons, we did not include units from the same tetrode from multiple sessions unless the tetrodes had been moved between those sessions. We further excluded a small number of neurons on the same tetrode that appeared to be potential duplicates based on waveforms and firing properties (e.g. firing rates, CV, and behavioral correlations), even though the tetrode had been moved. After recording was complete, we anesthetized rats and made small marker lesions by applying 5-10 µA current for 20 s on one or two wires of each tetrode. After perfusing the rats, tissue sections (at 40 µm) were stained with cresyl violet or with CD11b antibody and compared to the nearest atlas section (Paxinos, 2006).

## Data analysis

### Firing rates and neuron’s functional grouping

Firing rates were smoothed using a Gaussian kernel (30ms DS) and normalized (Z-scored) using the neuron’s session-wide mean and SD. Most analyses were done using this normalized firing rates except fraction of units (Fig. 2 (A),(F) bottom, using binned firing rates) and bursting analysis (Fig. 4, see below). Units are categorized into increase or decrease types using initiation score and into Contra>Ipsi or Ipsi>Contra types using selection score (statistically significant across trials, Wilcoxon signed rank test, *p*<0.05).

### Bursting

Spike bursts were detected using the Poisson surprise method (Legendy and Salcman 1985) with a surprise threshold of 5, and the burst ratio was calculated as the number of spikes fired in bursts divided by the number of all spikes in each unit. The CV of inter-spike intervals and burst ratio were calculated in each trial using a 3s time window before the *Go!* cue, and averaged across all trials (for sessions with/without proactive inhibition comparison) or across each trial type (for conditional differences).

### PCA analysis

was done largely as previously described (Gu *et al*., 2020). The smoothed, normalized average time series for contra and ipsi actions (500ms each, around Center Out, Supp. Fig. 2A) were used for PCA. This population activity matrix **R** is zero-centered, and after using MATALB ‘svd’ function, the PC scores (S) was calculated as S = **R**W, where W is the right singular vectors.

To match the analysis to the Initiation and Selection scores (Figure 2), we defined the Initiation Axis by connecting state space points 200ms before and at action initiation (averaging Contra and Ipsi actions), and the Selection Axis by connecting the mean of Contra trajectories to the mean of Ipsi trajectories (again using the epoch from -200 to 0ms relative to action initiation). The projections onto the Initiation and Selection Axis were calculated as the dot product of the state space position vector and the axis vector.

For mapping of before Go cue time series data (**R’**) into the contra and ipsi action related PC dimension, new PC scores were calculated by S’ = **R’**W after matching the zero values of **R’** to the matrix **R**. For subpopulation analysis (Fig 3. (B), (C)), selected unit’s population vector (e.g. **R**_(ipsi)_) and the right singular vectors (e.g. W_(ipsi)_) were used to calculate the selective unit’s PC scores (S_(ipsi)_ = **R**_(ipsi)_W_(ipsi)_).

### Permutation tests

To test if conditional differences are statistically significant, we ran permutation tests by randomly shuffling the two comparing trial conditions for each neuron (10000 shuffles). The original distance between the two comparing conditions were compared to the distance between shuffled trial conditions after projecting the distances onto the Initiation or Selection axis.

### Variability of neural trajectories

To examine how the increased variability of individual neurons affects population dynamics, we calculated the temporal variability of neural state-space trajectories. We included data from 3s time windows before the *Go!* cue, in each session (n=27) with at least 5 recorded units and a significant proactive inhibition effect. Trajectory variability V corresponds to the mean Euclidean distance between each time point of the trajectory and the center of the trajectory, i.e. how much the trajectory wanders around. For this analysis, we did not apply PCA to reduce dimensionality. Also, the trajectory variability was normalized by the number of neurons in each session. Thus, for example, if the normalized FR of population neurons (N) at time (t) is defined as a vector 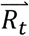, and the center of trajectory (mean FR across t=1…T, where T = 1500 for 3s time window with 500Hz sampling rate) is defined as a vector 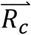Variability (V) of a neural trajectory is defined as:

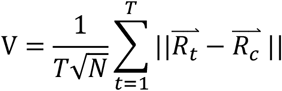

## Supplemental information

**Supplementary Figure 1**. (A) Brain slice examples of using CD11 antibody or Cresyl violet to mark electrode tip after making lesion. (B) Estimated locations of recorded cells, within coronal atlas sections (Paxinos & Watson 2006).

**Supplementary Figure 2**. Ipsi>Contra type cells are further divided into increase (A) and decrease (B) type cells. Both group of cells show significantly increase firing rates before *Go!* cues. Same formats as in the figure 2.

**Supplementary Figure 3**. (A) PCA was performed using 500 ms epoch around Center Out for contra and ipsi movements (averaged, normalized firing rates were concatenated). (B) Variance explained by each of the first 10 PCs. (C) The first 10 principal components. (D) Population dynamics of different trial outcomes (200ms around Go! cue) show different neural trajectory patterns. Permutation test shows positioning differences at *Go!* cue compared to correct contra go trials. The trials with wrong choice are biased toward ipsi action initiation at the time of *Go!* cue. Formats of permutation histograms are same as in figure 3.

**Supplementary Figure 4**. (A) Sessions with high CV (averaged across units) shows longer reaction times (mean reaction times of all trials). Each data point shows the value of each session. (B) Sessions with bigger increase of CV during proactive inhibition shows bigger proactive inhibition effects. (C) Sessions with proactive inhibition effect show bigger CV compared to the sessions without proactive inhibition effect in individual rats (one-tailed Wilcoxon signed rank test, *p*<0.05).

**Supplementary Figure 5**. Both Contra>Ipsi and Ipsi>Contra type of cells show increased CV and burst ratios with selective proactive inhibition, however only Contra>Ipsi cells shows a statistically significant effect (Wilcoxon signed rank test). Same format as in the figure 4D.

**Supplementary Figure 6**. Individual rat data shows firing rates (during 200ms before *Go!* cue) and CV (3s before *Go!* cue) differences between two conditions. *p<0.05 without correction,

**p<0.05 with Bonferroni correction (Wilcoxon signed rank tests in each subject). Error bar indicate ±S.E.M.

