## Supplementary figures and images for "Altered basal ganglia output during self-restraint"

### Supplementary Figures

#### *Supp. Figure 1*

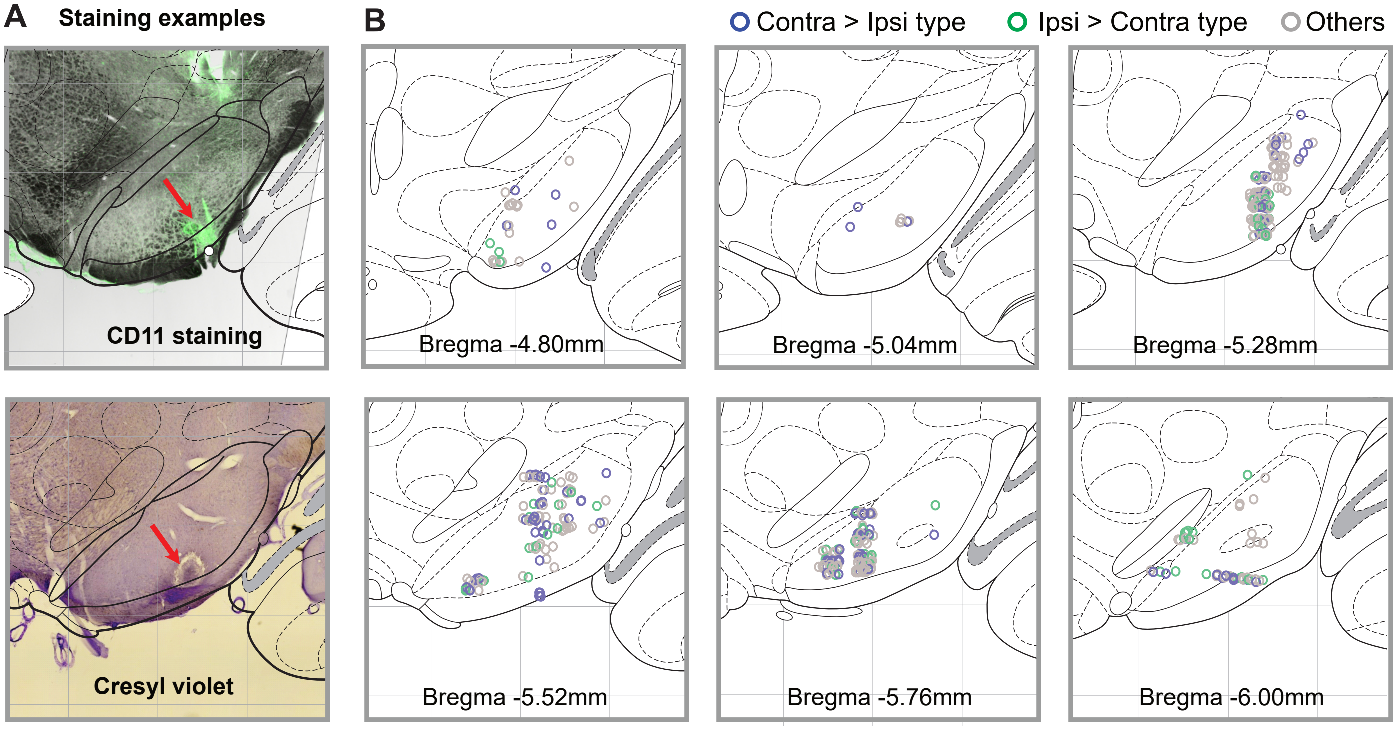


#### *Supp. Figure 2*

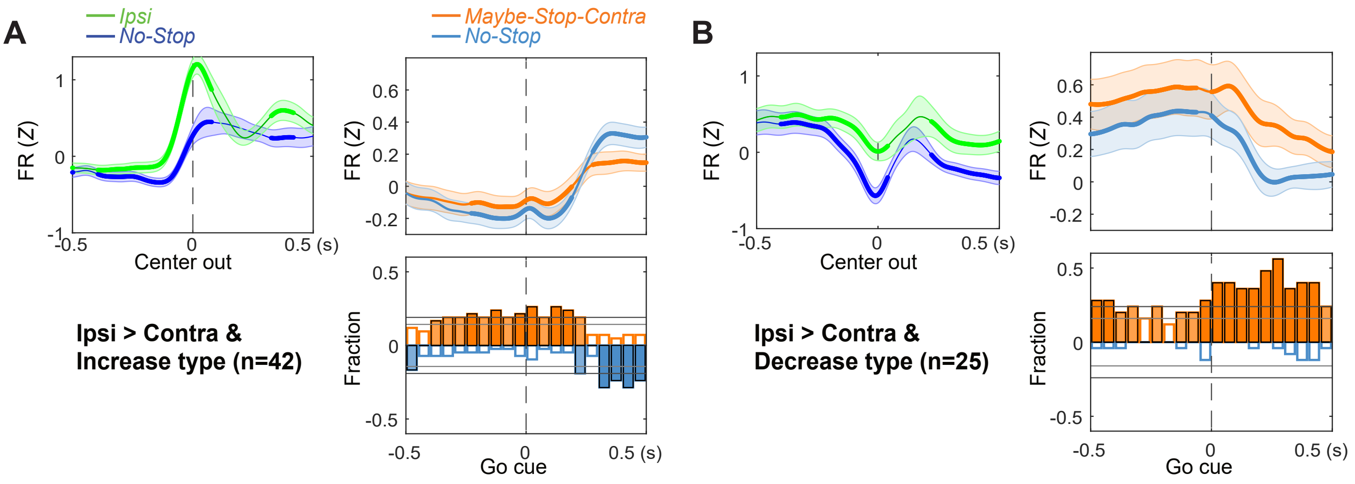

#### Supp. Figure 3

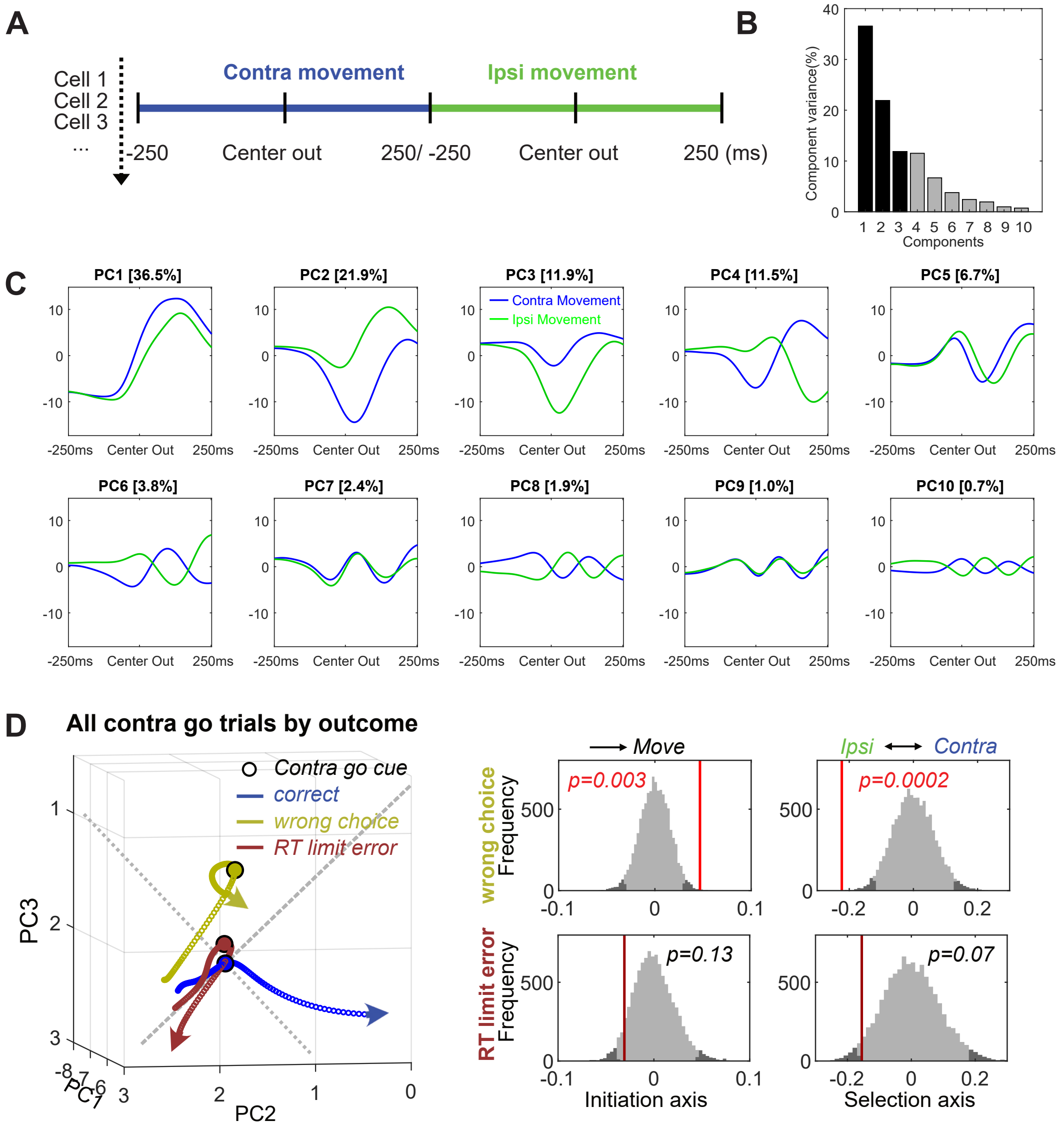

#### *Supp. Figure 4*

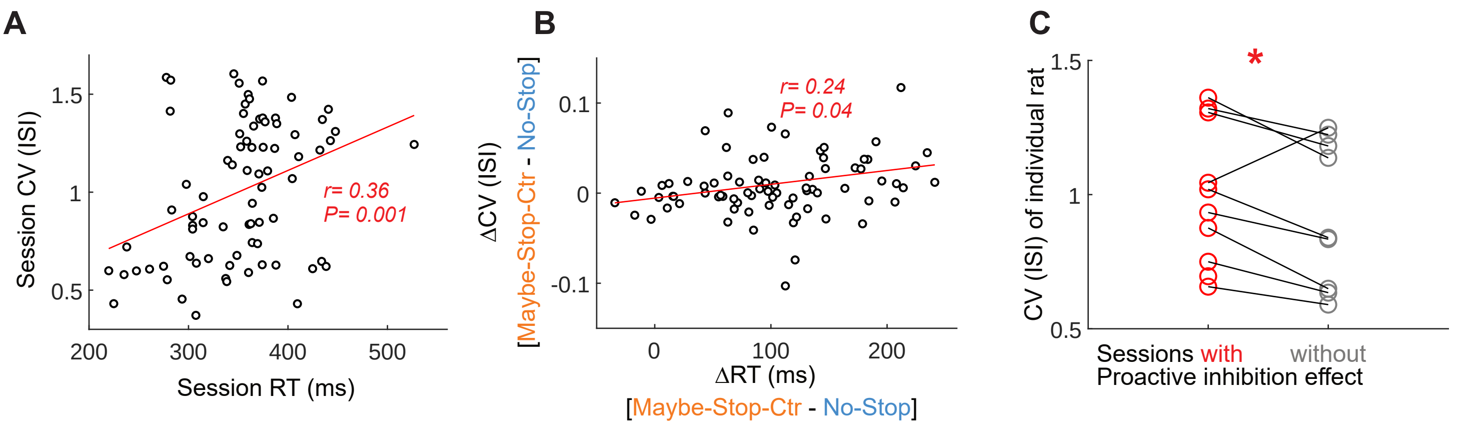


#### *Supp. Figure 5*

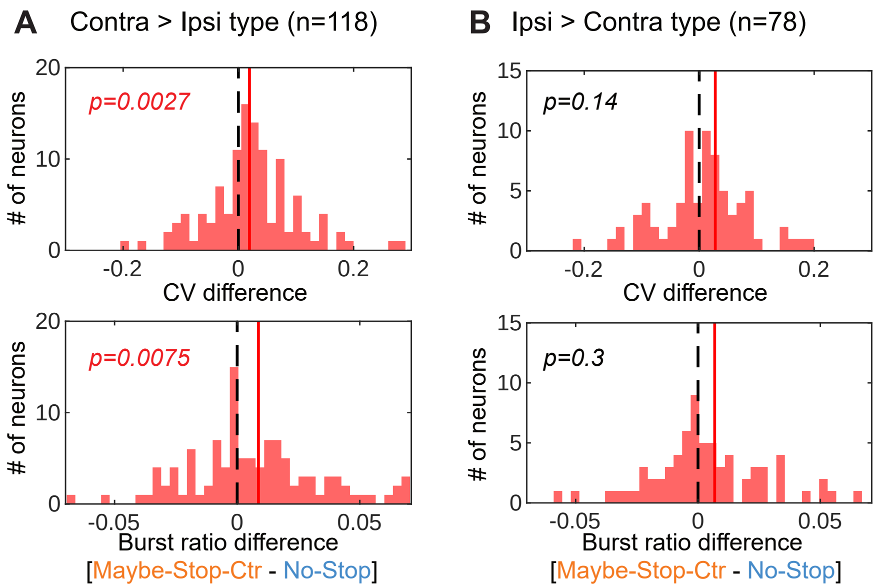


#### *Supp. Figure 6*

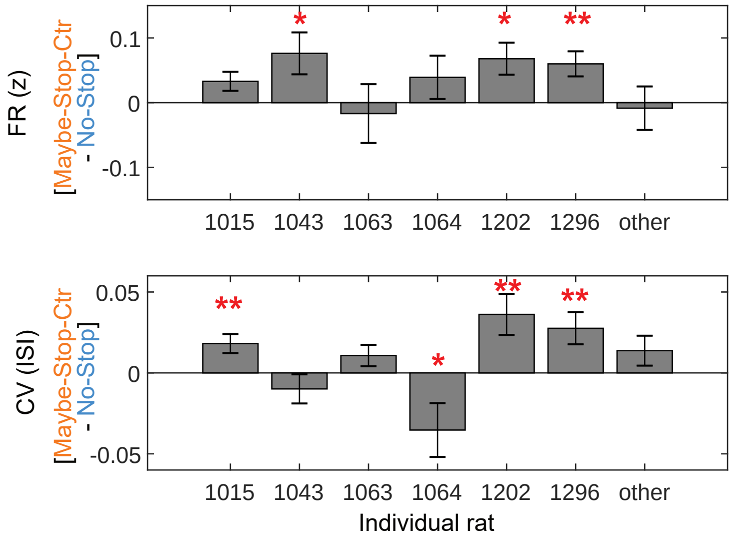
